# Efficient Genetic Code Expansion Without Host Genome Modifications

**DOI:** 10.1101/2024.05.15.594368

**Authors:** Alan Costello, Alexander A. Peterson, David L. Lanster, Zhiyi Li, Gavriela D. Carver, Ahmed H. Badran

**Affiliations:** Department of Chemistry The Scripps Research Institute; La Jolla, CA, 92037, USA; Department of Integrative Structural and Computational Biology The Scripps Research Institute; La Jolla, CA 92037, USA; Doctoral Program in Chemical and Biological Sciences The Scripps Research Institute; La Jolla, CA, 92037, USA; Department of Molecular Biology Princeton University, Princeton, NJ, 08544, USA

## Abstract

Supplementing translation with non-canonical amino acids (ncAAs) can yield protein sequences with new-to-nature functions, but existing ncAA incorporation strategies suffer from low efficiency and context dependence. We uncover codon usage as a previously unrecognized contributor to efficient genetic code expansion using non-native codons. Relying only on conventional *E. coli* strains with native ribosomes, we develop a novel plasmid-based codon compression strategy that minimizes context dependence and improves ncAA incorporation at quadruplet codons. We confirm that this strategy is compatible with all known genetic code expansion resources, which allows us to identify 12 mutually orthogonal tRNA–synthetase pairs. Enabled by these findings, we evolve and optimize five tRNA–synthetase pairs to incorporate a broad repertoire of ncAAs at orthogonal quadruplet codons. Finally, we extend these resources to an *in vivo* biosynthesis platform that can readily create >100 new-to-nature peptide macrocycles bearing up to three unique ncAAs. Given the generality of our approach and streamlined resources, our findings will accelerate innovations in multiplexed genetic code expansion and enable the discovery of chemically diverse biomolecules for researcher-defined applications.

## Introduction

Genetic code expansion – where non-canonical amino acids (ncAAs) are added to the central dogma – has provided foundational tools to study and manipulate biological processes^1^. Pioneering labs have shown that genetic code expansion requires two components: heterologous tRNA–synthetase pairs engineered to accept researcher-defined ncAAs, and “blank” codons available for assignment to ncAAs during translation. Historically, the codon of choice has been the amber (UAG) stop, and considerable effort has focused on improving UAG-dependent ncAA incorporation through strain and ribosome engineering^2–7^. Innovations in genome synthesis and unnatural nucleobases have provided new sequences that can be used for multiplexed ncAA incorporation^8–10^. Despite these developments, *in vivo* genetic code expansion often suffers from low ncAA incorporation efficiencies and context dependence for unknown reasons. Conversely, *in vitro* genetic code expansion can access chemically diversified peptides through precise control over tRNA aminoacylation and abundance^11^, but can result in low protein yield and require extensive tRNA preparation. Combining the *in vitro* chemical diversity with the scalability and throughput of *in vivo* of genetic code expansion could streamline the discovery of bioactive peptides and non-canonical polymers.

While many aspects of translation (e.g., codon sequence, tRNA aminoacylation, and ribosome components) have been investigated to address these limitations and improve ncAA incorporation outcomes^12–20^, the mRNA has received comparatively little attention. We set out to investigate the contribution of mRNA sequence to ncAA incorporation using quadruplet codon translation. Quadruplet-decoding tRNAs (qtRNAs) can “read” a four-base codon complementary to an engineered four-base anticodon, resulting in a +1 frameshift and restoration of the open reading frame during translation^21^. While quadruplet codon translation offers new codons for multiplexed ncAA incorporation, it is less efficient than triplet decoding^19^. Nonetheless, quadruplet decoding is compatible with native ribosomes since 1) qtRNAs are active in *in vitro* translation^22^, 2) qtRNAs can be evolved to improve their *in vivo* activities^19,23^, and 3) aminoacylation by engineered synthetases can occur in an anticodon-independent manner^24^. We hypothesized that two mRNA parameters may impact quadruplet codon translation. First, codon usage in natural genes can fine-tune the rate^25^, fidelity^26^, and efficiency^27^ of translation. While amino acid^28,29^ identities near UAG can affect ncAA incorporation^30,31^, the impact of neighboring codons on quadruplet decoding remains unknown. Second, the unavoidable sequence overlap between triplet and quadruplet codons may sequester qtRNAs from their intended decoding events and inadvertently promote frameshift errors. In agreement with this, the most commonly used qtRNAs decode sequences that overlap with low-usage triplet codons^19,23,32^, yet no analysis of qtRNA-dependent mistranslation has been reported to date.

Here, we explored the impact of mRNA codon usage on quadruplet decoding, finding a preference for a high-usage triplet codon immediately downstream of a quadruplet codon. To improve on-target decoding and eliminate off-target activities in the same gene, we generalized this observation such that all plasmid-borne genes were recoded using a single representative codon for each of the 20 canonical amino acids (cAAs). We optimized virtually all parameters predicted to affect qtRNA-dependent decoding using this framework, including plasmid designs, copy numbers, promoter strengths, qtRNA sequences, and synthetase identities. Our improved resources catalyze the efficient incorporation of multiple unique ncAAs into proteins without requiring host *E. coli* cell alteration. Finally, we applied this approach towards the biosynthesis of peptide macrocycles encoding up to three unique ncAAs in living cells for the first time, bridging the gap between *in vitro* and *in vivo* genetic code expansion approaches. Our simple recoding strategy was refined into a user-friendly “plug-and-play” technology for expansive quadruplet codon translation.

## Results

### Local and Remote Codons Influence Observable Quadruplet Decoding

Local sequence context and codon usage can influence natural translation and UAG decoding^27,30,31,33^. To explore its impact on quadruplet decoding, we used a superfolder GFP (sfGFP) reporter where the permissive Y151 was replaced by a quadruplet codon^4^, and varied the codon usage of five upstream and downstream residues (**Figure 1a**). Using cAA- and ncAA-specific qtRNAs^19,32,34^, we sorted cells based on high vs. low sfGFP expression (**Extended Data Figure 1**). We repeated this analysis using a UAG codon at Y151 to ensure that our findings are specific to quadruplet decoding. Sorted clones showed a wider sfGFP distribution when decoding quadruplet codons vs. UAG, suggesting that quadruplet decoding is more sensitive to local sequence contexts (**Extended Data Figure 4a, b**). Next-generation sequencing (NGS) showed little change in codon preference between UAG bins (**Figure 1b**, **Extended Data Figure 4c**), whereas quadruplet decoding populations showed clear codon usage biases (**Extended Data Figure 4d-f**). Specifically, high sfGFP bins consistently showed a strong preference for a high usage codon immediately 3’ to the quadruplet codon (i.e., +1 position; **Figure 1c-e**). We also observed codon bias at other positions within our libraries, namely at the –4, –1, +2, and +3 positions with respect to the decoding site. However, these positions did not show a recurring trend between UAGA vs. AGGA codons, or between quadruplet vs. triplet decoding. We hypothesized that these observations may therefore reflect context-dependent effects and were not explored further. We validated our NGS results by assaying the top sequences from the Top 5% and Top 1% pools (**Figure 1f**), and confirmed amino acid incorporation using purified sfGFP from the naïve library and all sorted pools using mass spectrometry (**Extended Data Figure 2-3**). Finally, we tested high vs. low usage codon substitutions at the +1 position when decoding AGGA at other positions in sfGFP, finding that +1 recoding to high usage codons is either beneficial or of no consequence (**Figure 1g**).

**Figure 1.**
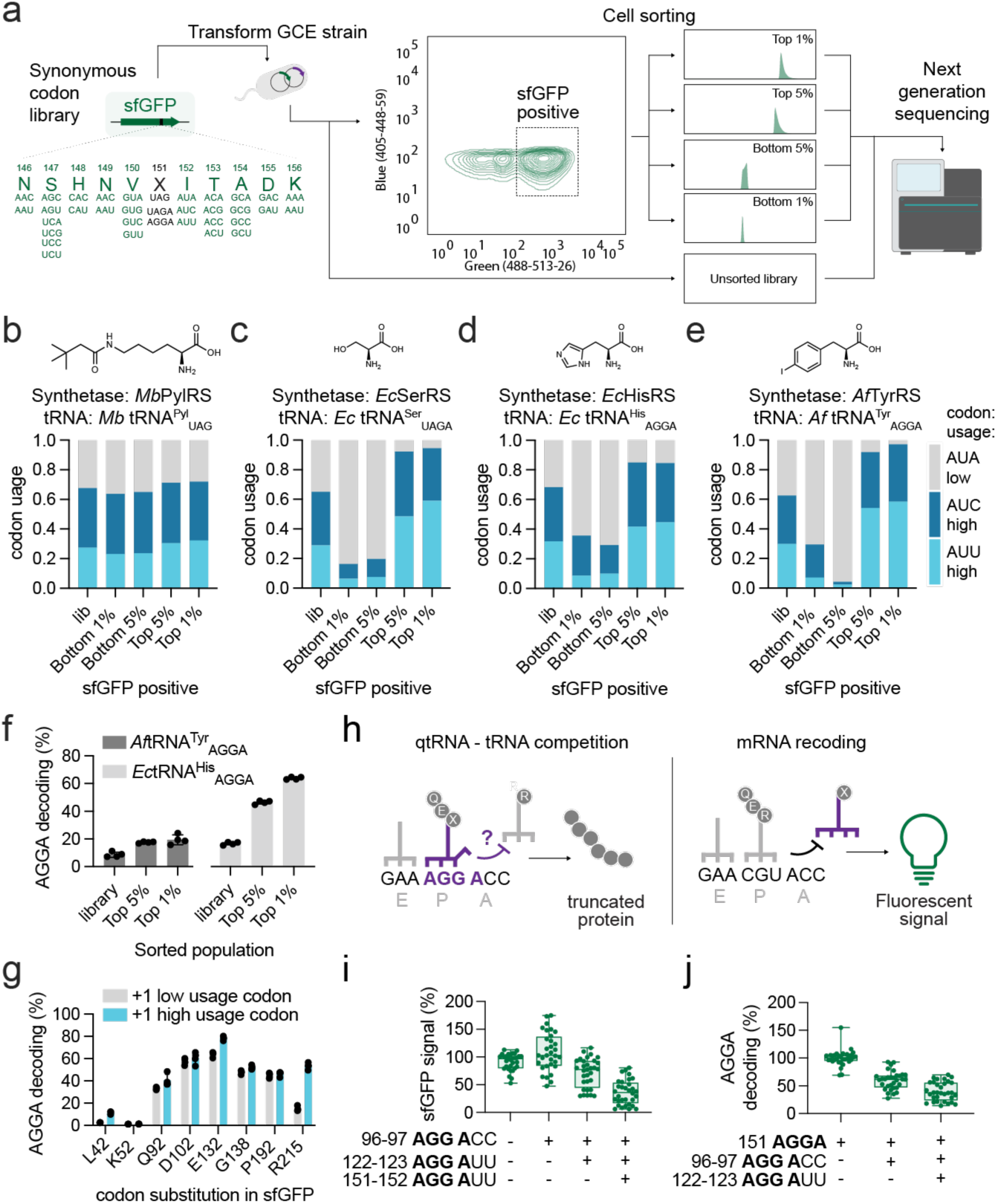
| Local and Remote Codons Influence Apparent Quadruplet Decoding. **a)** Library design and screening approach to investigate the influence of codon usage on quadruplet decoding. Codon abundance before and after fluorescence activated cell sorting (FACS) is shown at the +1 position. I152 is encoded by three codons AUA, AUC, and AUU. AUA is a low usage codon while AUC and AUU are both high usage codons in *E. coli*. The codon abundance in transcripts at the +1 (I152) position is shown for: **b)** *Mb* tRNA^Pyl^_UAG_ with *Mb*PylRS incorporating N6-(tert-butoxycarbonyl)-lysine (BocK); **c)** *Ec* tRNA^Ser^(evo2)_UAGA_ incorporating serine; **d)** *Ec* tRNA^His^_AGGA_ incorporating histidine; **e)** *Af* tRNA^Tyr^(A01)_AGGA_ with *Af*TyrRS(G5) incorporating para-iodophenylalanine (pIF). **f)** Variants from the top 1% and 5% populations showed greater AGGA decoding than the library average using *Af*TyrRS(G5)-*Af* tRNA^Tyr^(A01)_AGGA_ and *Ec* tRNA^His^_AGGA_ (n = 8). **g)** Testing the influence of low vs. high usage codons at the +1-position using multiple positions in sfGFP and *Ec* tRNA^His^_AGGA_ (n = 4). **h)** Schematic representation of putative competition between a qtRNA and tRNA whose codons overlap at nucleotides 1-3. Recoding the mRNA to remove triplet codons that overlap with quadruplet codons eliminates this competition at unintended sites. **i)** Using an all-triplet codon sfGFP reporter, introduction of in-frame AGG codons at positions R96, R122, and Y151 results in mistranslation by many different AGGA-decoding qtRNAs and reduced apparent GFP translation (n = 7). **j)** Apparent AGGA decoding at Y151 is similarly reduced when AGG codons are introduced in-frame at positions R96 and R122, resulting in reduced apparent GFP translation (n = 7). Wherever reported, AGGA decoding % was calculated by normalizing OD_600_-corrected sfGFP fluorescence values to a wild-type sfGFP control.

Next, we speculated that qtRNAs likely do not distinguish between on-target (which corrects the translation frame) and off-target decoding (which creates *de novo* frameshift mutations during translation). Since quadruplet codons often include low usage triplet codons at the first three positions^19^, we hypothesized that restricting mRNAs to high usage codons would eliminate off-target decoding and improve protein purity (**Figure 1h**). We recoded our reporter genes, antibiotic resistance markers, and plasmid replication proteins to use a single high usage codon for each amino acid. We expected this recoding may impair protein function in some cases but did not observe any issues with maintenance or growth of strains carrying recoded plasmids. Using different AGGA-decoding qtRNAs, we observed robust quadruplet decoding in our optimized circuit architecture lacking AGG triplet codons (**Extended Data Figure 5a, b**), whereas reintroducing AGG in sfGFP reduced apparent decoding efficiency through presumed off-target frameshifting (**Figure 1i-j**, **Extended Data Figure 5c**). These results collectively implicate codon usage as an unrecognized yet important determinant of efficient, on-target quadruplet decoding.

### Engineered tRNA–Synthetase Pairs are Active in Recoded Genetic Circuits

Introducing multiple unique ncAAs into a single protein requires mutually orthogonal tRNA– synthetase pairs^35^. To ensure that our recoding strategy can be extended to multiplexing studies, we first synthesized a *Methanosarcina barkeri* (*Mb*)-derived PylRS–tRNA^Pyl^_CUA_ using our high usage codon set and evaluated its activity using a sensitized UAG luciferase reporter (**Figure 2a**, **Extended Data Figure 6a**)^19,36,37^. We optimized synthetase plasmid copy number, induction conditions, and tRNA expression (**Extended Data Figure 6b-d**), finding that our recoded components performed as effectively as their *E. coli* codon-optimized counterparts (**Figure 2b**). Aiming to nominate the most robust components for multiplexed ncAA incorporation, we next evaluated representatives of all tRNA–synthetase pairs reported in *E. coli* to date using UAG decoding (for both cAA and ncAA incorporation)^12,15,32,37–49^. In some cases, we tested related synthetases with differing ncAA scopes, altered protein expression to improve host tolerance, and optimized tRNA decoding efficiencies and specificities (**Extended Data Figure 7-10**).

**Figure 2.**
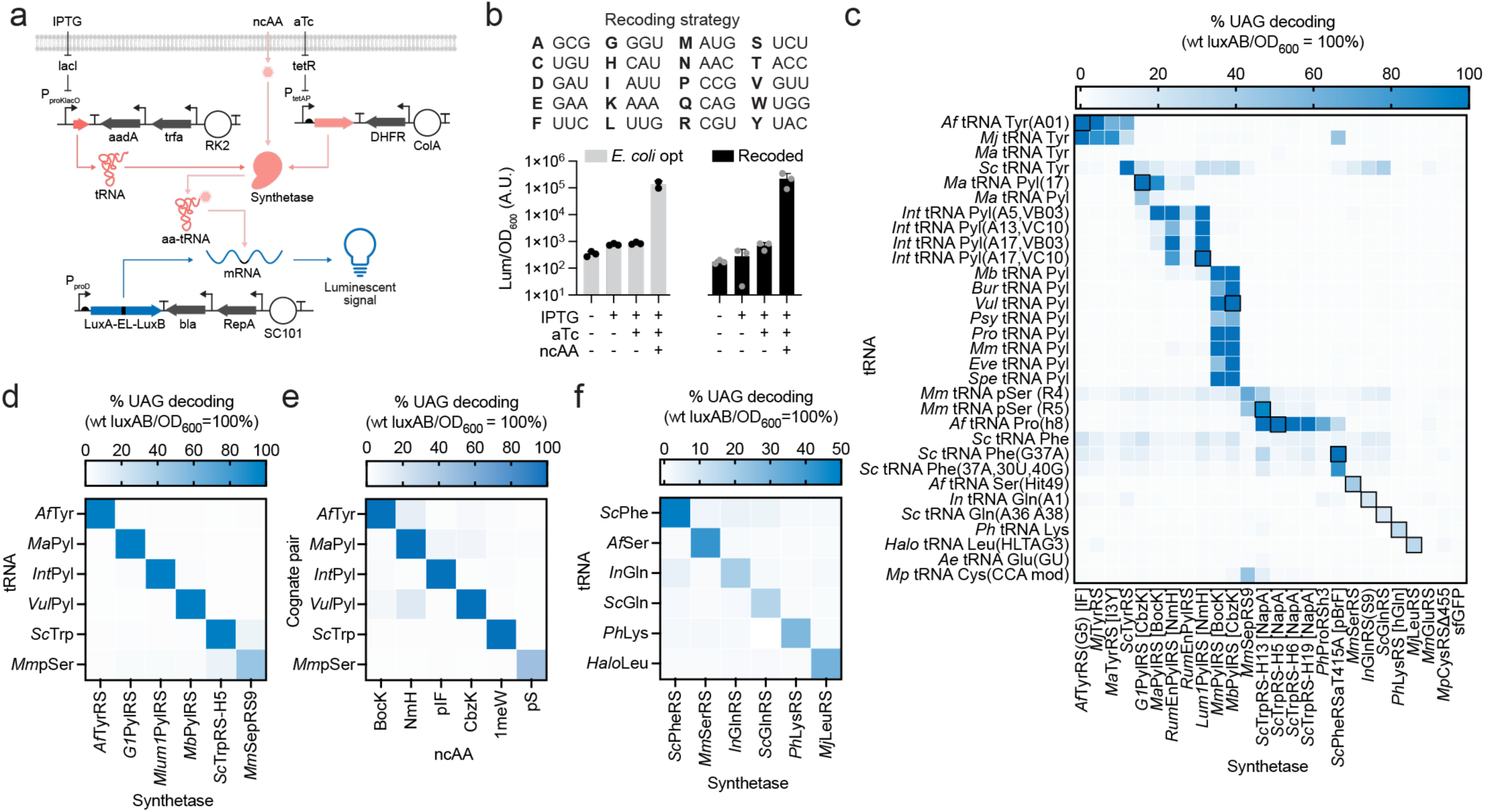
| Twelve Mutually Orthogonal tRNA–Synthetase Pairs and Validation in Recoded Circuits. **a)** Schematic representation of the tRNA–synthetase benchmarking assay. All tRNAs are expressed from the IPTG-controlled promoter (P_proK_–lacO) and synthetase proteins are produced from the aTc-responsive promoter (P_tetA_). The LuxAB reporter genes are separated by an engineered linker (EL) that contains an amber (UAG) codon. Origins of replication (white circles) and antibiotic resistance genes (gray arrows) are shown. **b)** All plasmid-encoded protein-coding genes were recoded to reflect a single high usage triplet codon for each canonical amino acid as shown. The *Mb*PylRS – *Mb*tRNA^Pyl^_UAG_ was tested using this recoding strategy and compared to conventional *E. coli* codon optimized genes. Luminescence values are normalized to OD_600_ and raw arbitrary units (A.U.) are shown (n=3). **c)** Heterologous synthetase (x-axis) and tRNA (y-axis) genes were combinatorially tested using the UAG LuxAB reporter. All ncAAs shown in parentheses on the x-axis were included in growth media at 1mM each, whereas *Mm*SepRS was tested alongside EF-Sep and *Tk*SerK. This analysis yielded a priority list of 12 mutually orthogonal tRNA–synthetase pairs (black boxes). Reporter activity is normalized to wild type LuxAB expression in all cases (n = 4). **d)** Validation of mutual orthogonality of the six identified ncAA-specific tRNA–synthetase pairs from the primary screen (n = 4). **e)** Validation of low crosstalk between ncAAs used by all six tRNA–synthetase pairs: N6-(tert-butoxycarbonyl)-lysine (BocK), N-methylhistidine (NmH), para-iodophenylalanine (pIF), N6-((benzyloxy)carbonyl)-lysine (CbzK), 1-methyltryptophan (1meW), and phosphoserine (pS). All ncAAs were included at 1mM each, whereas *Mm*SepRS was tested alongside EF-Sep and *Tk*SerK (n = 4). **f)** Validation of mutual orthogonality of the six identified cAA-specific tRNA–synthetase pairs from the primary screen (n = 4). Wherever reported, UAG decoding % was calculated by normalizing OD-corrected luminescence values to a wild-type LuxAB control.

We assayed 25 representative synthetases alongside 32 cognate tRNAs (800 combinations) and quantified UAG decoding as a percentage of a wild-type luciferase reporter (**Figure 2c**, see **Methods**). We uncovered 69 tRNA–synthetase pairs with >20% decoding efficiency, which could be assigned to 12 mutually orthogonal clusters. Clusters 1-6 are ncAA-specific and 7-12 are cAA-specific (**Figure 2d-f**). Cluster 1 contains TyrRS–tRNA^Tyr^_UAG_ pairs: *Af*TryRS, *Mj*TyrRS, *Ma*TyrRS, and *Sc*TyrRS. Clusters 2-4 contain orthogonal PylRS–tRNA^Pyl^_UAG_ variants^39^: *Ma*PylRS, *G1*PylRS, *Rum*PylRS, *Mlum1*RS, *Mm*PylRS, and *Mb*PylRS. Clusters 1-4 represent the current state-of-the-art for mutually orthogonal tRNA–synthetase pairs in *E. coli*^32^. Cluster 5 contains *Sc*TrpRS with a new *Sc*tRNA^Trp^(M13)_UAG_ we developed (**Extended Data Figure 8**). Cluster 6 contained two *Mm*SepRS– tRNA^Sep^_UAG_ variants^45,46^, which required the L-serine kinase *Tk*SerK^50^ and elongation factor EF-Sep^45^ for phosphoserine incorporation (**Extended Data Figure 9**). Clusters 7-12 correspond to 6 cAA-incorporating tRNA–synthetase pairs that may serve as starting points for further evolution: *Sc*PheRS^44^, *Mm*SerRS^49^, *In*GlnRS^43^, *Sc*GlnRS^43^, *Ph*LysRS^12^, and *Mj*LeuRS^47^ (**Figure 2f**). In addition to being the largest analysis of its kind to date, this dataset uncovered 12 mutually orthogonal tRNA– synthetase pairs and confirmed that codon compression is compatible with cAA and ncAA incorporation.

### qtRNA Directed Evolution Improves Quadruplet Decoding

Multiplexed ncAA incorporation will require engineered qtRNAs that efficiently decode non-overlapping quadruplet codons^19,32^. Having established five mutually orthogonal tRNA–synthetase pairs for UAG decoding and ncAA incorporation, we next optimized their quadruplet decoding activities. The tRNA–synthetase pairs *G1*PylRS, *Af*TyrRS, *Lum1*RS, and *Mm*PylRS have been engineered to decode quadruplet codons^32^, although their decoding efficiencies ranged from ∼0-20% in our assays (**Extended Data Figure 10a**). Reducing plasmid burden by condensing all components from three to two plasmids improved efficiencies to 8-100% depending on the tRNA–synthetase pair, albeit with high background activity in some cases (**Extended Data Figure 10b**). These poor activities may restrict future multiplexing studies. Whereas the above ncAA qtRNAs were created by swapping anticodon stem loop segments from established variants^51^, we have shown that directed evolution can significantly improve qtRNA decoding efficiencies^19^.

We used established chloramphenicol acyltransferase (CAT) positive selections and barnase negative selections^52^ to improve qtRNA activities (**Figure 3a**). We systematically targeted the qtRNA anticodon stem loop, acceptor stem, Ψ-arm, and/or D-arm with site saturation mutagenesis and subjected the libraries to iterative positive-negative selection campaigns. These efforts improved the qtRNA–synthetase pairs described above to decode orthogonal quadruplet codons with up to 100% efficiency in a two-plasmid system: *G1*PylRS (3.4-fold improvement in dynamic range using AUAG), *Af*TyrRS (1.6-fold; CUAG), *Lum1*RS (5.3-fold; AGGA), and *Mm*PylRS (9.8-fold; UAGA) (**Figure 3b-e**, **Extended Data Figure 11-14, Extended Data Table 1-4**). Seeking to supplement our ncAA repertoire with tryptophan analogs, we further developed CGGA as a new quadruplet codon for the *Sc*TrpRS–tRNA^Trp^(M13) pair (**Extended Data Figure 15**) and used directed evolution to discover *Sc*tRNA^Trp^_CGGA_ variants with up to a 38.1-fold dynamic range in the three-plasmid circuit and 5-fold dynamic range in the two-plasmid system (**Figure 3f**, **g, Extended Data Table 5**). We confirmed these enhancements in quadruplet decoding between the starting and evolved qtRNAs through western blot (**Extended Data Figure 16**) and mass spectrometry analyses (**Extended Data Figure 17**). Cumulatively, these campaigns yielded five ncAA-incorporating, mutually orthogonal qtRNA– synthetase pairs that effectively function in living cells.

**Figure 3.**
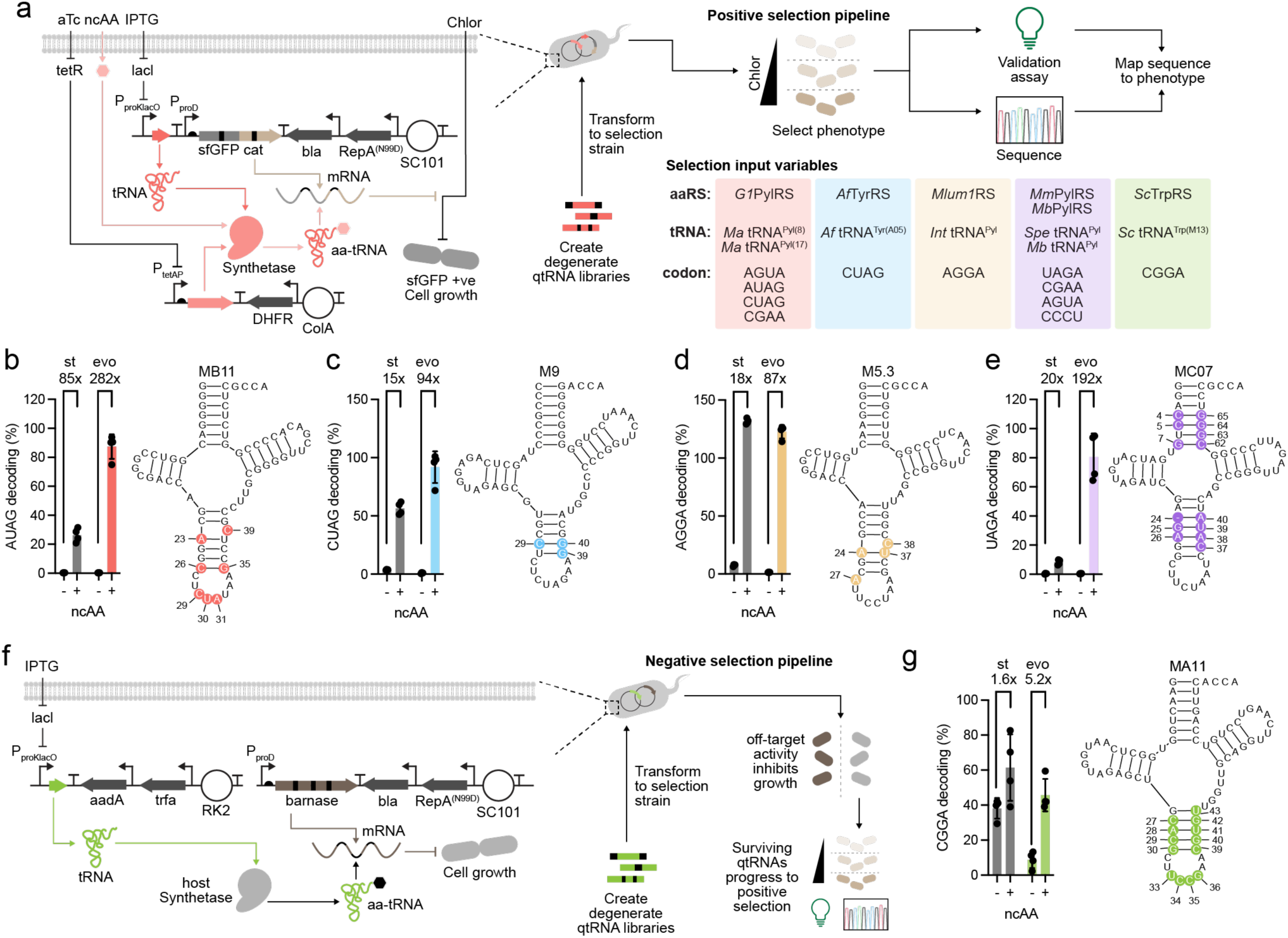
| Directed Evolution of Highly Efficient qtRNAs for ncAA Incorporation. **a)** Schematic representation of the selection strategy to evolve improved qtRNA activities in the recoded framework: qtRNA were subjected to positive selection by decoding quadruplet codons in an sfGFP–chloramphenicol acetyltransferase (cat) fusion gene. Starting (st) and evolved (evo) qtRNA genes were validated by decoding their cognate quadruplet codon at position Y151 in sfGFP. In each case, quadruplet decoding is reported for the final evolved variant and the discovered mutations are mapped onto the starting qtRNA scaffold: **b)** *Ma*tRNA^Pyl^_AUAG_ evolved to mutant (MB11), aminoacylated by *G1*PylRS (n = 4); **c)** *Af*tRNA^Tyr^_CUAG_ evolved to mutant (M9), aminoacylated by *Af*TyrRS (n = 4); **d)** *Int*tRNA^Pyl^_AGGA_ evolved to mutant (M5.3), aminoacylated by *Mlum1PylRS* (n = 4); **e)** *Spe*tRNA^Pyl^_UAGA_ evolved to mutant (MC07), aminoacylated by *Mm*PylRS (n = 4); **f)** In the case of *Sc*tRNA^Trp^_CGGA_, a negative selection was first implemented to eliminate ncAA-independent aminoacylation by host synthetases, then subjected to positive selection. **g)** *Sc*tRNA^Trp^_CGGA_ evolved to mutant (MA11), aminoacylated by *Sc*TrpRS (n = 4). Wherever reported, quadruplet decoding % was calculated by normalizing OD-corrected fluorescence values to a wild-type sfGFP control.

### Optimizing Synthetase ncAA Scope and qtRNA Expression

Multiplex ncAA incorporation would benefit from synthetases with a broad substrate scope to enable the introduction of a large repertoire of ncAAs using only a handful of mutually orthogonal synthetase-tRNA pairs. Since synthetases are often reported to incorporate only a single ncAA, we screened active site variants and homologs of our prioritized tRNA–synthetase pairs for broadened ncAA scope beyond their cognate substrates (**Extended Data Figure 18-22, Extended Data Table 6-11**). We prioritized synthetases that can access at least three ncAAs with >10% efficiency. These analyses uncovered variants of our five prioritized synthetases that together provide access to 47 unique ncAAs with diverse physicochemical properties (**Figure 4a**).

**Figure 4.**
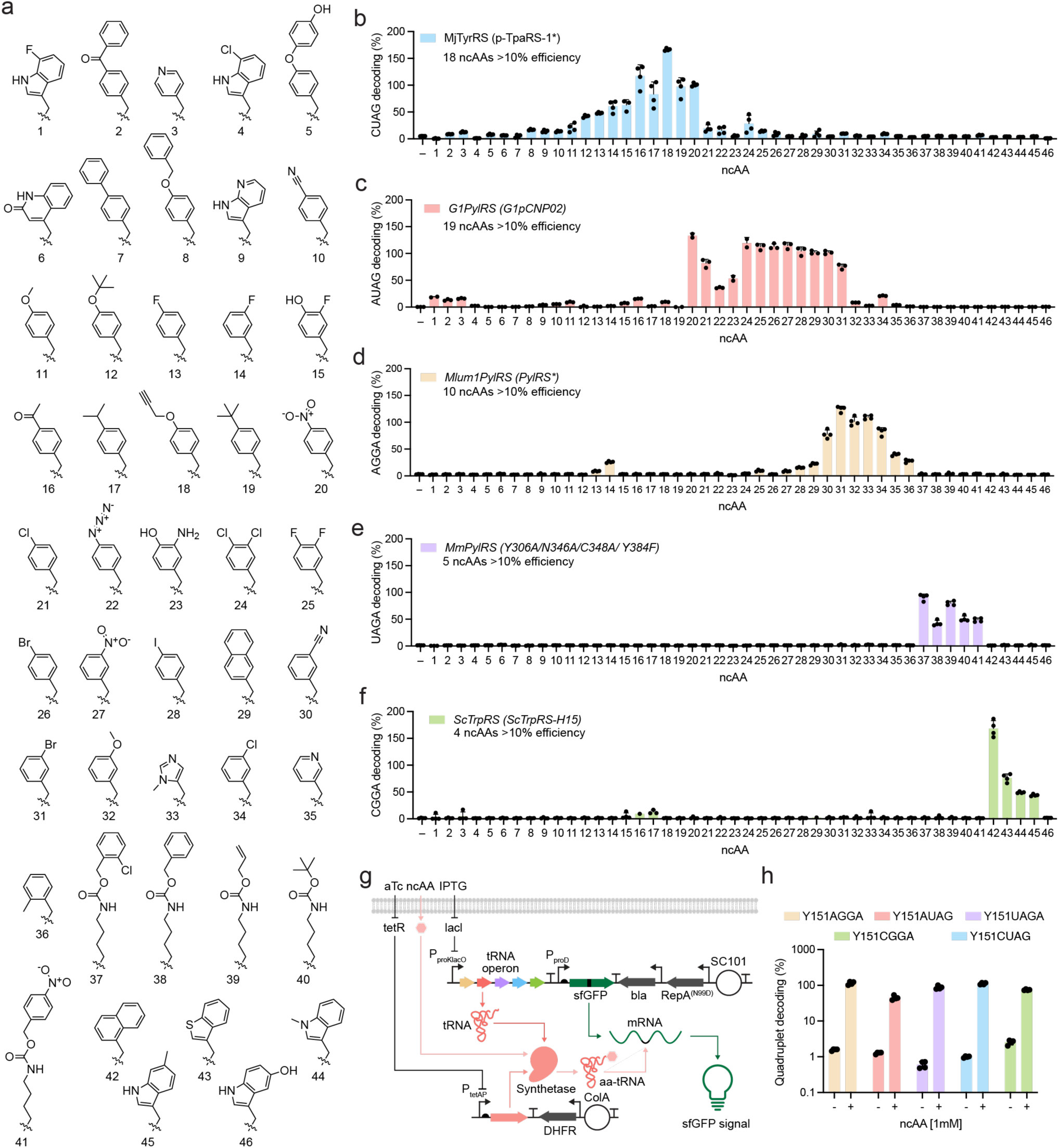
| Validation of Synthetases with Broad, Non-Overlapping ncAA Substrate Scope. **a)** Prioritized ncAAs for incorporation at quadruplet codons using optimized qtRNA–synthetase pairs. All shown ncAAs showed detectable signal using an sfGFP reporter using at least one qtRNA–synthetase pair. A detailed list of tested synthetase variants (**Extended Data Tables 6-10**) and ncAAs (**Extended Data Table 11**) are provided. The ncAA substrate scope is shown for the representative synthetases: **b)** *Mj*TyrRS (p-TpaRS-1*) **c)** *G1*PylRS (G1pCNP02) **d)** *Mlum1*PylRS (PylHRS*) **e)** *Mm*PylRS (Y306A/N346A/C348A/ Y384F) **f)** *Sc*TrpRS (ScTrpRS-H15) **g)** Schematic representation of a multicistronic qtRNA operon and testing using dedicated synthetases by decoding quadruplet codons at Y151 in sfGFP. **h)** Decoding efficiency of sfGFP Y151 mutated to a quadruplet codon for each evolved qtRNA encoded on an optimized multicistronic operon. Decoding was evaluated in the absence (–) and presence (+) of a cognate ncAA at 1mM. Wherever reported, quadruplet decoding % was calculated by normalizing OD-corrected fluorescence values to a wild-type sfGFP control.

Briefly, *Mj*TyrRS (set 1 = Y32I, L65I, Q109M, D158G, L162V, V164G)^15^ was developed to incorporate *para*-(2-tetrazole)phenylalanine, and fortuitously uses diverse *para*-substituted and *para*-/*meta*-disubstituted phenylalanine derivatives (**Figure 4b**). We note that this *Mj*TyrRS also carried mutations that improve tolerance to qtRNA anticodon changes (Y230K, C231K, P232K, H283Q, D286S)^53^. *G1*PylRS (L124S, Y125F, Y204W, A221S, W237Y)^54^ was developed to incorporate cyanopyridylalanine, but is capable of charging a large repertoire of related *meta*- and *para*-substituted phenylalanine and tyrosine derivatives (**Figure 4c**). M*lum1*RS (L125I, Y126F, M129G, V168F, Y205F) was grafted from a reported *Mb*PylRS^55^ that uses histidine derivatives, which allows Lum1RS to charge derivatives of histidine, thiophene, naphthalene, as well as *meta*- or *ortho*-substituted phenylalanines (**Figure 4d**). *Mm*PylRS (Y271A, Y349F)^56^ was developed to incorporate lysine analogs, which we reproduce here (**Figure 4e**). *Sc*TrpRS (Y106V, E141P, T233C, I253C)^57^ was evolved to use 6-methyltryptophan, and we show that it can also tolerate related analogs (**Figure 4f**).

Finally, we optimized qtRNA production from a multicistronic cassette since this can improve multiplexed quadruplet decoding efficiency^19,32^. We used a reported 4-qtRNA cassette and mined the *E. coli* genome for inter-tRNA sequences (20-60 nucleotides) to use as spacers for our 5^th^ qtRNA (**Figure 4g**, **Extended Data Table 2**). Multiple spacers resulted in expression of the 5^th^ qtRNA and robust quadruplet decoding (**Extended Data Figure 23**). The best qtRNA expression cassette resulted in excellent quadruplet decoding for all five optimized qtRNAs in a ncAA-dependent manner (**Figure 4h**). Together, these chosen synthetases and qtRNA expression constructs enable efficient decoding of quadruplet codons and incorporation of diverse ncAAs in conventional *E. coli* cells.

### Robust Biosynthesis of Non-Natural Macrocycles Using Quadruplet Decoding

Strategies to introduce ncAAs into peptide macrocycles have yielded natural product-inspired molecules^9,58^ and probes of protein function^11,59,60^. Advances in *in vitro* tRNA aminoacylation and cell-free translation have catalyzed the discovery of chemically diverse bioactive macrocycles using mRNA display, usually enriched through binding to purified proteins^61,62^. While recent advances in genome rewriting have enabled the biosynthesis of macrocycles encoding two ncAAs or hydroxy acids, these methods are restricted to a single engineered bacterium and require *in vitro* cyclization of a cell-generated thioester intermediate^9,10^. Efforts to create chemically complex macrocycle libraries *in vivo* remain limited by poor ncAA incorporation and codon decoding efficiencies. Addressing these limitations would enable streamlined macrocycle discovery *in vivo* by increasing the chemical repertoire available to protein translation and potentially capitalizing on directed evolution paradigms to optimize attributes beyond binding.

We explored an intracellular biosynthesis strategy to generate macrocyclic peptides that encode diverse, researcher-assigned ncAAs. We adapted a reported split-intein circular ligation of peptides and proteins (SICLOPPS) system using the *Nostoc punctiforme* (Npu)^63^ engineered to create cyclo-CLLFVY^64^, and combined it with our optimized system for ncAA incorporation (**Figure 5a**). We first confirmed that active Npu generated cyclo-CLLFVY with robust signal, whereas an active site mutant (C1A)^65^ showed no detectable macrocycle production by LC-MS (**Figure 5b**). Npu plasmid copy number and cyclic peptide sequence both influenced product yield (**Extended Data Figure 24**), in line with prior observations^66^. Next, we extended this approach to ncAA incorporation using quadruplet decoding in Npu-derived macrocycles. We co-transformed strains with each of the five optimized qtRNA–synthetase pairs and Npu genes encoding quadruplet codons at variable macrocycle positions, finding macrocycle products with robust ion counts comparable to cyclo-CLLFVY (**Figure 5c**, **Extended Data File 1**). In all cases, we observe the correct primary mass at M+1. In these assays, incorporation of 3-bromophenylalanine (3BrF) was a key validation step since bromine has a unique isotopic pattern (M and M+3 in roughly equal abundance) and is otherwise poorly abundant in *E. coli*. We observed this isotopic pattern for a 3BrF-containing macrocycle, providing confidence in the correct assignment of the target macrocycle from a heterogenous sample (**Extended Data Figure 25**).

**Figure 5.**
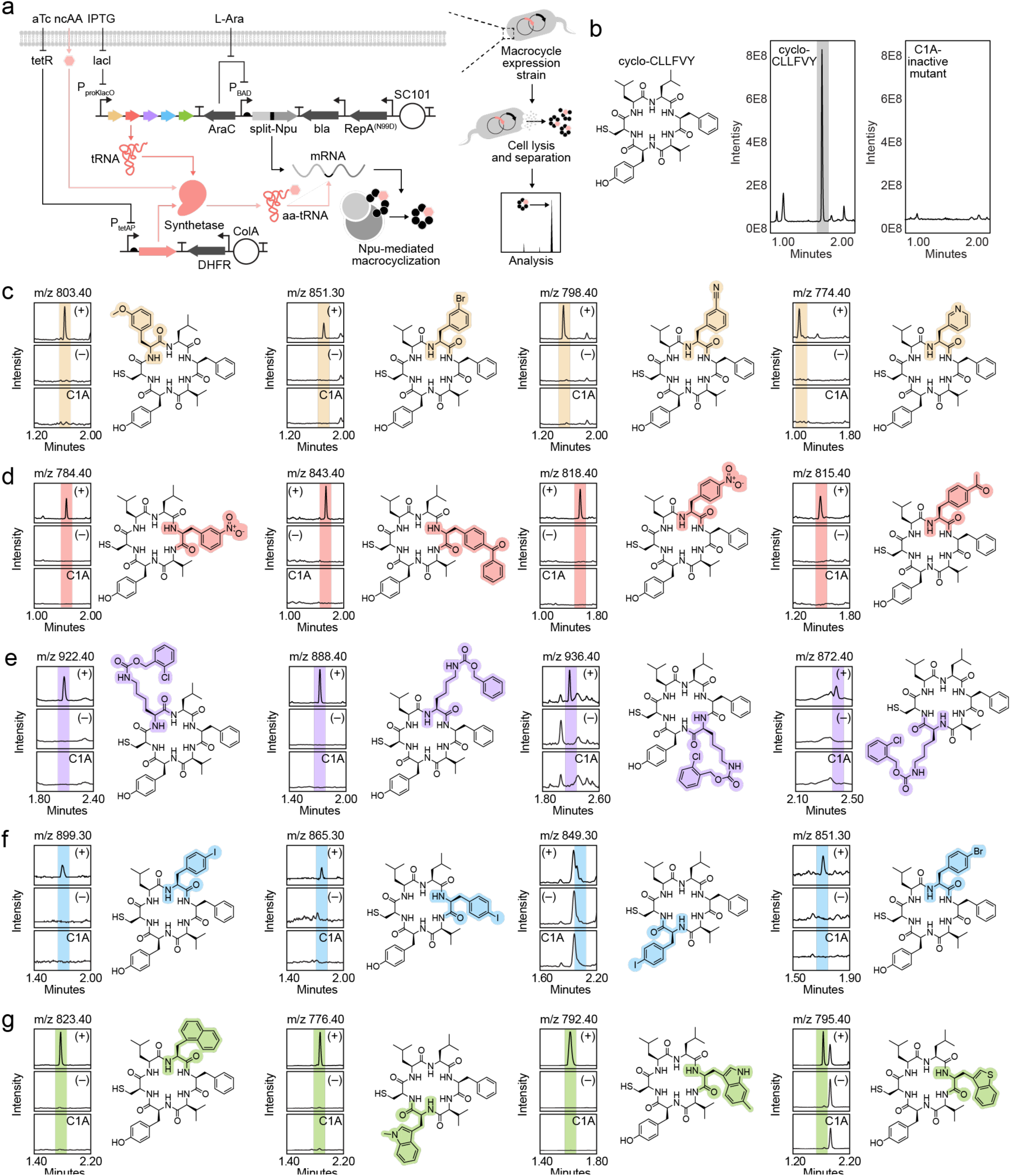
| Intracellular Biosynthesis of ncAA-Encoding Peptide Macrocycles Through Optimized Quadruplet Decoding. **a)** Schematic representation of the macrocycle biosynthesis pipeline. The split-*Npu* genes are expressed from an arabinose-controlled pBAD promoter, whereas the qtRNAs and synthetase are expressed from IPTG- and aTc-inducible promoters, respectively. Desired macrocycle sequences (with or without quadruplet codons) are incorporated between the split-Npu genes, resulting in splicing and macrocyclization after translation. Following expression, cells were lysed, and organic soluble components isolated for analysis by LC-MS. **b)** The model macrocycle cyclo-CLLFVY is robustly biosynthesized by Npu, whereas the C1A active site mutation ablates macrocycle generation. This sequence was used as a template for all subsequent ncAA incorporation studies. In parts **c-g**, individual quadruplet codons were tested at positions 2-6 of cyclo-CLLFVY (cysteine at is required for intein-mediated peptide macrocyclization) and combined with the cognate synthetase plasmids to explore scope and positional tolerance for ncAA incorporation. Examples of extracted ion chromatogram (XIC) at the anticipated mass +H are shown (50 unique macrocycles in **Extended Data File 3**). In each case, XICs for the indicated mass are shown for the following conditions: + ncAA (+), – ncAA (–), and C1A inactivated intein + ncAA (C1A). Color highlights for the ncAAs indicate the used qtRNA–synthetase pairs consistent with Figure 4: **c)** AGGA decoding in yellow by *Int* tRNA^Pyl^_AGGA_ M5.3 and *Mlum1*PylRS (PylHRS*); **d)** AUAG decoding in red by *Ma* tRNA^Pyl^_AUAG_ MB11 and *G1*PylRS (G1pCNP02); **e)** UAGA decoding in purple by *Spe* tRNA^Pyl^_UAGA_ MC07 and *Mm*PylRS (Y306A/N346A/C348A/Y384F); **f)** CUAG decoding in blue by *Af* tRNA^Tyr^_CUAG_ M9 and *Mj*TyrRS (p-TpaRS-1*); and **g)** CGGA decoding in green by *Sc* tRNA^Trp^_CGGA_ MA11 and *Sc*TrpRS (ScTrpRS-H15).

We note that apparent variability in macrocycle extracted ion chromatograms (XIC) may be attributed to splicing, ionization, and/or protonation differences and cannot be directly compared. To provide insight into yield and corroborate our peak assignment, we compared our LC-MS profiles to authentic standards generated using solid-phase peptide synthesis (SPPS), which showed identical XIC retention times and confirmed correct peak assignment. By serially titrating each synthetic macrocycle, we could estimate cellular yield of select macrocycles. Across the five synthetase pairs, we observe up to 2 µM macrocycle yields following extraction (**Extended Data Figure 26, Extended Data File 4**). In total, we generated 50 macrocycles encoding a single unique ncAA each, highlighting the scope of our optimized qtRNA–synthetase pairs.

### Designer Non-Natural Macrocycles that Incorporate Multiple Unique ncAAs

Prior work by our lab and others has shown that multiplexing quadruplet decoding events can often lead to poor overall protein production^19,32^. Bolstered by robust decoding efficiencies and broad ncAA scope in model macrocycles, we tested multiplexed incorporation of distinct ncAAs through quadruplet codon decoding. To our knowledge, no strategy has demonstrated the cellular biosynthesis of macrocycles with more than two distinct ncAAs or hydroxy acids. We first explored the incorporation of a single ncAA multiple times within the same macrocycle, finding that our approach resulted in excellent production of macrocycles containing up to three identical ncAAs (**Figure 6a-b**, **Supplementary Data File 5**).

**Figure 6.**
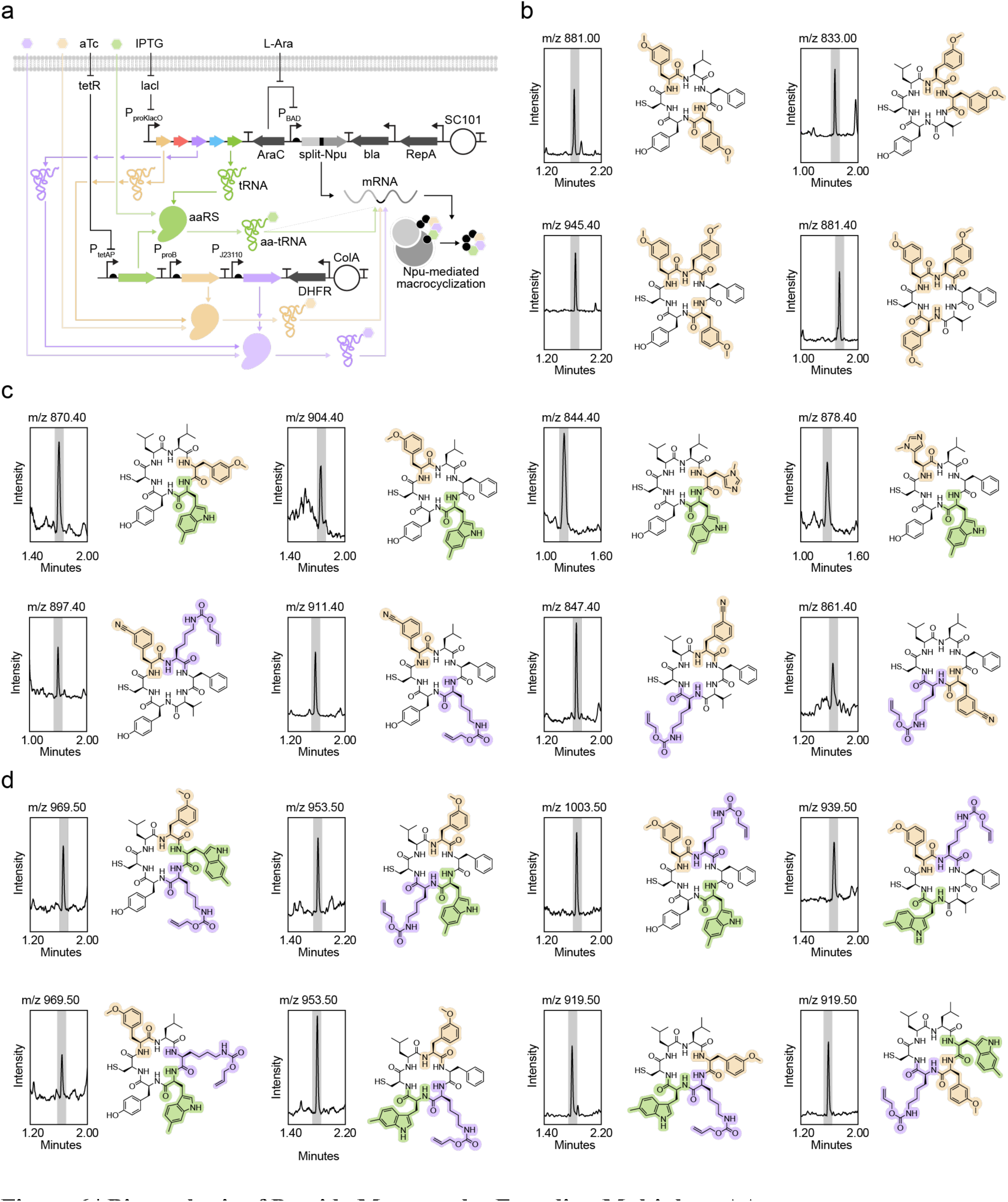
| Biosynthesis of Peptide Macrocycles Encoding Multiple ncAAs. **a)** Schematic representation of the macrocycle biosynthesis pipeline for multiplexed ncAA incorporation. The inducible promoters and extraction strategy are identical to Figure 5. **b)** Demonstration of processive ncAA incorporation into the model macrocycle cyclo-CLLFVY using 3-methoxy-L-phenylalanine. Extracted ion chromatogram (XIC) at the anticipated mass +H are shown. **c)** Incorporation of two unique ncAAs in cyclo-CLLFVY using combinations of AGGA+CGGA or AGGA+UAGA codons. Extracted ion chromatogram (XIC) at the anticipated mass +H are shown. In total, 36 macrocycles encoding two unique ncAAs were generated through quadruplet decoding (**Extended Data File 5**). **d)** Incorporation of three unique ncAAs in cyclo-CLLFVY using combinations of AGGA+CGGA+UAGA codons. Extracted ion chromatogram (XIC) at the anticipated mass +H are shown. In total, 17 macrocycles encoding three unique ncAA were generated through quadruplet decoding (**Extended Data File 6**). HRMS analysis for all samples is detailed in (**Extended Data Table 13**).

We then designed and validated genetic constructs to co-express two or three of our prioritized synthetases (*Mlum1*RS + *Mm*PylRS, *Mlum1*RS + *Sc*TrpRS, or *Mlum1*RS + *Mm*PylRS + *Sc*TrpRS; **Figure 6a**, **Extended Data Figure 27**). We substituted codons in the model macrocycle cyclo-CLLFVY with combinations of UAGA, CGGA, and/or AGGA, and monitored macrocycle biosynthesis by LC-MS. Using these components, we generated 72 unique macrocycles that encoded two or three ncAAs without extensive protein or macrocycle purification (**Figure 6b-c**, **Extended Data Figure 28, Supplementary Data File 5**). While poor processivity in decoding adjacent quadruplet codons can reduce protein yield^67,68^, our evolved qtRNAs show robust processivity as evidenced by the biosynthesis of 17 macrocycles carrying three consecutive ncAAs. In summary, we showcase – for the first time – optimized resources for genetic code expansion capable of generating new-to-nature peptide macrocycles containing up to three distinct ncAA residues.

## Discussion

Genetic code expansion was pioneered to enable the synthesis of biopolymers with diverse, researcher-defined chemical sequences beyond nature. Notably, natural peptide macrocycles can have activities that are desirable in the biotechnology sector, including: antibiotics^69^, immunosuppressants^70^, and antitumor compounds^71,72^. Natural peptide macrocycles are generated by non-ribosomal peptide synthetases (NRPS) or through post-translational modifications of ribosomally-translated peptides^73,74^. Engineering these biosynthetic pathways to create new products can be difficult since the responsible enzymes are often recalcitrant to changes that faithfully alter substrate scope. In addition, optimizing the expression of such large enzyme complexes in heterologous hosts can be challenging. While traditional solid-phase peptide synthesis strategies can create novel structures, they lack the means to readily refine peptide macrocycle bioactivities. Technologies that access chemically diversified peptide macrocycles while capitalizing on the genetically encodable nature of protein translation would offer new opportunities to discover and refine their bioactivities, capitalizing on mature directed evolution approaches or emerging machine learning-based methods. Despite considerable progress, strategies to effectively biosynthesize peptides or proteins bearing new chemistries in a programmable manner in living cells remains an outstanding challenge.

Aiming to bridge this gap, genetic code expansion was pioneered by the Schultz lab as a means to alter the physicochemical properties of proteins in living cells^75^. While the linkage between a tRNA and the ncAA would later be addressed through the directed evolution of dedicated aminoacyl tRNA synthetases^76^, the availability of unassigned codons has consistently been a key limitation^77^. Recent innovations in genome synthesis^78,79^ and artificial nucleobases^80^ have begun to address this fundamental issue by erasing redundant codons from the host genome or by creating *de novo* codons, respectively. Yet genome synthesis requires a deep understanding of regulatory signatures embedded within genes, and the use of artificial nucleobases relies on expertise in their synthesis and activation for use as substrates *in vivo*. While quadruplet decoding may address these issues, the corresponding codons are thought to be poorly read by the cell’s translation machinery.

To realize the promise of genetic code expansion, our lab and others have recently developed new biotechnologies to improve qtRNAs, synthetases, and the ribosome in high throughput^34,36,68,81^ . Strategies that enhance the incorporation of diverse ncAAs will catalyze innovations in protein design and bioengineering that go beyond the constraints of the natural proteinogenic amino acids. In this report, we investigated how an underexplored parameter – mRNA codon composition – can influence this process. We find that local and global mRNA codon usage can have a significant impact on quadruplet decoding and ncAA incorporation efficiency. We develop new genetic circuits that obviate these issues, improve qtRNA decoding and expression, and optimize synthetase dosing and ncAA scope. Together, our optimized resources were distilled into a streamlined plasmid series (**Extended Data Table 14**) that can incorporate up to three ncAAs across various protein contexts.

Our approach here aims to combine the codon flexibility and chemical scope of *in vitro* macrocycle synthesis with the scalability and throughput of *in vivo* genetic code expansion. Accordingly, it provides key strengths beyond state-of-the-art methods. First, the chemical breadth provided by five mutually orthogonal qtRNA–synthetase pairs allows access to diverse macrocycles using a single DNA sequence. Future macrocycle discovery efforts may therefore benefit from smaller DNA libraries that can be easily elaborated using unique ncAA combinations, overcoming transformation efficiency bottlenecks that often hinder directed evolution campaigns. Second, our strategy does not require any host genome modifications to enable multiplexed ncAA incorporation, which will allow researchers to explore biosynthetic strategies using ncAAs in their favored *E. coli* strains. Third, our combined macrocycle biosynthesis and ncAA incorporation strategy is fully genetically encodable and does not require purification of any intermediates from cells to complete peptide cyclization. Our approach can therefore leverage design–build–test-learn cycles to accelerate small molecule discovery through macrocycle-dependent selections. Taken together, we envision that these favorable properties will catalyze new innovations using *in vivo* genetic code expansion to build chemically diverse polymers in a programmable manner.

## Methods

### General Methods

All DNA manipulations were performed using NEB Turbo cells (New England Biolabs) or Mach1F cells, which are Mach1 T1^R^ cells (Thermo Fisher Scientific) mated with S2057 F’ to constitutively provide TetR and LacI^81^. All DNA manipulations were performed using NEB Turbo cells (New England Biolabs) or Mach1F cells, which are Mach1 T1^R^ cells (Thermo Fisher Scientific) mated with S2057 F’ to constitutively provide TetR and LacI^81^. All assays were performed with *E. coli* S3489 as previously described^82^. Water was purified using a MilliQ water purification system (Millipore). All amplifications were carried out using Phusion U Hot Start DNA Polymerase (Life Technologies) or repliQa HiFi ToughMix (Quantabio). MinElute PCR Purification Kit (Qiagen) was used to purify all PCR products to 10 μl final volume, which was quantified using a NanoDrop 1000 Spectrophotometer (Thermo Fisher Scientific). All plasmids were constructed by USER (Uracil-Specific Excision Reagent; Endonuclease VIII and Uracil-DNA Glycosylase) cloning^83^.

### USER Cloning and Sequencing

A single internal deoxyuracil base was included at 10–20 bases from the 5′ end of each primer used to amplify plasmid components. This region is described as the USER junction, which specifies the homology required for correct assembly. USER junctions were designed to contain minimal secondary structure, have 42 °C < *T*_m_ < 70 °C, and begin with a deoxyadenosine and end with a deoxythymine (to be replaced by deoxyuridine). For USER assembly, an equimolar ratio (up to 1 pmol each) of PCR products carrying complementary USER junctions were mixed in a 10 μl reaction containing 0.75 units DpnI (New England Biolabs), 0.75 units USER enzyme (New England Biolabs), 1 μl of CutSmart Buffer (50 mM potassium acetate, 20 mM Tris-acetate, 10 mM magnesium acetate, 100 μg mL^−1^ BSA at pH 7.9; New England Biolabs). The reactions were incubated at 37 °C for 20-45 min, followed by heating at 80 °C for 2 mins and slow cooling to 12 °C at 0.1 °C s^−1^ in a thermocycler. The hybridized constructs were directly used for heat-shock transformation of chemically competent NEB Turbo *E. coli* cells or Mach1F *E. coli* cells. 2xYT (United States Biological) agar plates (1.8%) supplemented with the appropriate antibiotic(s) were used to select for transformants. In all cases, cloned plasmids were verified by Sanger sequencing using template generated by the TempliPhi 500 Amplification Kit (GE Life Sciences) according to the manufacturers protocol or by Nanopore sequencing of purified plasmid DNA (Primordium Labs). For high-throughput sequencing of synonymous codon library pools, plasmid DNA was harvested from sorted populations using conventional silica-based column purification. The region of interest was amplified, barcoded, and sequenced using a MiSeq instrument (Illumina). For data analysis, a 33bp or 34 bp insert between forward and reverse primers was extracted and the frequencies of silent codons was quantified. Data was filtered for only synonymous mutations to eliminate issues that may result from incorrect synthesis of the library primers. During our synonymous codon library generation efforts, we noted that creating a for serine at the –4 position required six codons, but this could not be exclusively introduced using a degenerate codon approach. In this case, we elected to use the degenerate codon WSN (where W=A/T, S=C/G, and N=A/C/G/T), which includes all 6 serine codons, as well as codons corresponding to arginine, threonine, cysteine, and tryptophan. In our next-generation sequence (NGS) analysis, we excluded the reads which did not match the intended amino acid sequence (i.e. serine) as our goal was to only identify codon usage biases which affect quadruplet decoding and not local amino acid contexts.

### Chemically Competent Cell Preparation and Transformation

An overnight culture was diluted 100-fold in 2xYT media containing maintenance antibiotics and grown at 37 °C with shaking at 350 rpm to OD_600_ = 0.5–0.7. Cells were pelleted by centrifugation at 5000 × rcf for 5 min at 0°C. Supernatant was decanted, and the pellet was resuspended in the residual media, keeping it on ice for 10-15 min and gently shaking at short regular intervals. TSS (2xYT medium supplemented with 5% v/v DMSO, 10% w/v PEG 3350, and 20 mM MgCl_2_) was added to the resuspend cells and mixed by gently swirling at a volume of 10% of the original culture. The cell suspension was then aliquoted, flash-frozen in liquid nitrogen, and stored at -80°C until needed. To transform cells, 100-400 μL aliquots of competent cells were thawed from –80 °C on ice for 15 min. An equal volume of KCM solution (100 mM KCl, 30 mM CaCl_2_, 50 mM MgCl_2_) was added to the tube and mixed gently by tapping the tube. Plasmid DNA or USER reactions were mixed with TSS/KCM/cell mixtures and left on ice for 10-30 min. Then cell/DNA mixtures were heat-shocked at 42°C for 90 seconds. The transformation was then chilled on ice for 2 min, then added to 1 mL of 2xYT media. Cells were allowed to recover at 37 °C with shaking at 350 RPM for 1h before plating on 2xYT agar (1.5%) containing the appropriate antibiotics and incubated at 37 °C for 16−18 hours.

### Luminescence Assays

Early log-phase (OD_600_ = 0.3-0.5) S3489 cells carrying the reporter plasmid (RP) grown in 2xYT (United States Biological) were made chemically competent, transformed with the desired tRNA–synthetase expression plasmids (EPs), and recovered for 2h in 2xYT (United States Biological). Transformations were plated on 1.8% agar-2xYT plates (United States Biological) supplemented with carbenicillin (15 µg mL^−1^), trimethoprim (3 µg mL^−1^), and spectinomycin (30 µg mL^−1^). The plates were incubated for 12–18 h in a 37 °C incubator. Colonies transformed with the appropriate EPs were picked the following day and grown in Davis Rich Media (DRM) containing carbenicillin (15 µg mL^−1^), trimethoprim (3 µg mL^−1^), and spectinomycin (30 µg mL^−1^) for 18 h. Following overnight growth of the EP/RP-carrying strains, cultures were diluted 250-fold into fresh DRM supplemented with carbenicillin (15 µg mL^−1^), trimethoprim (3 µg mL^−1^), and spectinomycin (30 µg mL^−1^). The cultures were induced with anhydrotetracycline (aTc) (100 ng mL^−1^), IPTG (1 mM), and supplemented with relevant ncAAs (1 mM). After 4-6 h of growth in a 37°C incubator shaking at 900 rpm, 150 μL of culture was transferred to a 96-well black wall, clear bottom plate (Costar) and measured for OD_600_ and luminescence. Each plasmid combination was assayed in 4 biological replicates. Luminescence activities were tabulated at OD_600_ = 0.1-0.5 in all cases. Where luminescent signal is reported as a % of wild type reporter activity, a control RP encoding a wild-type luciferase was grown in parallel and used for normalization across assays.

### Fluorescence Assays

Chemically competent S3489 cells were transformed with combinations of tRNA–synthetase EPs alongside sfGFP RPs. In all cases, transformations were recovered for 2h in 2xYT (United States Biological), then plated on 1.8% agar-2xYT plates (United States Biological) supplemented with carbenicillin (50 µg mL^−1^) and trimethoprim (10 µg mL^−1^) for one- or two-plasmid circuits, or carbenicillin (15 µg mL^−1^), trimethoprim (3 µg mL^−1^), and spectinomycin (30 µg mL^−1^) for three-plasmid circuits. Plates were incubated for 12–18 h in a 37 °C incubator. Colonies were picked the following day and grown in DRM containing appropriate antibiotics. After growth for 16-24h at 37°C shaking at 900 rpm, cultures were diluted 100- to 500-fold in DRM with antibiotics, inducers, and ncAAs as above. Following another 16-24h at 37°C shaking at 900 rpm, 150 μL of culture was transferred to a 96-well black wall, clear bottom plate (Costar) and measured for OD_600_ and fluorescence values. The reporter protein sfGFP was recorded by excitation at 485 nm and emission at 510 nm. In all cases, values were recorded using an Infinite M1000 Pro microplate reader (Tecan) or Spark plate reader (Tecan). Each variant was assayed in 4-8 biological replicates. Fluorescent protein yields were normalized to culture OD_600_ in all cases. Where sfGFP yield is reported as a % of wild type sfGFP, a control RP encoding a wild type sfGFP was grown in parallel and used for normalization across assays.

### qtRNA Directed Evolution

qtRNA libraries were made by PCR with degenerate base primers to amplify the qtRNA of interest. Libraries were assembled by USER cloning (see above) and transformed into S3489 cells carrying an aaRS expression plasmid. Library transformations were recovered for 1 h in DRM with aaRS and tRNA induction, and where applicable cognate ncAA (1mM). Transformations were plated by spreading on DRM agar chloramphenicol selection plates (below) and grown at 37 °C for up to 96 h. DRM plates: DRM media was combined 4:1 with a 5% molten agar solution. All maintenance antibiotics were added to the DRM agar: carbenicillin (50 µg mL^−1^), trimethoprim (10 µg mL^−1^), along with aaRS and qtRNA induction: aTc (100ng mL^-1^) and IPTG (1 mM). For positive selections, cognate ncAA (1mM) was added as a powder to molten DRM agar and mixed by stir bar. A titration of chloramphenicol from 32-0 (µg mL^-1^) was made in molten DRM agar with all required small molecules. To validate colony resistance to chloramphenicol, colonies found to grow above the strain MCI were picked into DRM with maintenance antibiotics overnight. These cultures were used to inoculate liquid culture titration of chloramphenicol to validate the phenotype.

### Protein Purification and Analysis

S3489 strains transformed with synthetase expression plasmids and reporter plasmids (sfGFP with C-terminal His-tag) were grown with maintenance antibiotics, aTc 100 (ng mL^-1^), IPTG (1 mM), and where relevant ncAA (1 mM) in 4 mL of DRM. Overnight cultures were pelleted, and supernatant removed. To each pellet, 0.5 mL BugBuster (Millipore) was added and incubated for 30 min at room temperature with gentle rocking. Soluble protein was separated by centrifugation at 16,000 × g for 20 min and recovering supernatant (soluble total protein lysate). 300 µL of soluble lysate was loaded onto a His-Spin Protein Mini-prep column (Zymo) and purified using the manufacturer’s protocol with one change: centrifugation steps were done at 700 x g for 2 min. All samples were eluted in 150 µL of elution buffer. Soluble protein lysate or nickel-NTA-purified protein was visualized by mixing 10µg of lysate or 5µL of eluted protein from purification with 4 µL NuPAGE LDS Sample Buffer (Invitrogen), 10 mM dithiothreitol (DTT), and dH_2_O up to 16 µL, and run on a 4-12% Bis-Tris NuPAGE gel (Invitrogen).

### Immunoblotting

Protein material was resolved by SDS-polyacrylamide gel electrophoresis (SDS-PAGE) in 17-well 4–12% Bis-tris NuPAGE gels (Invitrogen) in 1 x MES-SDS buffer (Invitrogen). Samples were transferred to nitrocellulose (NC) membrane (Amersham Protran) using a semi-dry transfer apparatus (BioRad). Membranes were blocked for 1 h at room temperature with StartingBlock (Thermo Scientific) in phosphate-buffered saline (PBS). Membranes were incubated with primary antibody (c-Myc Monoclonal Antibody 9E10) (Thermo Fisher) at a 1:1000 dilution in TBST with 5% bovine serum albumin (BSA), overnight with shaking at 4 °C. Membranes were exposed to fluorophore-conjugated secondary antibodies (IRDye® 680RD Donkey anti-Mouse IgG Secondary Antibody) (Li-Cor) diluted 1:5000 in TBST with 5% BSA at room temperature. Signals were recorded with a Li-Cor fluorescence imager using a ChemiDoc instrument (BioRad).

### TOF-MS Analysis

Salt concentration of Ni-NTA-purified sfGFP purified products was reduced in samples by buffer exchange with Amicon (Millipore) column centrifugation. Protein samples were then subjected to LC-MS analysis. The analysis was performed on an LC-MS consisting of a Waters I-Class LC and a Waters Xevo G2-XS TOF which uses a LeuEnk lockmass and is calibrated against sodium formate clusters. An Acquity BEH C4 column (1.7 µm, 2.1×55 mm) was used with a 5-99% B gradient lasting 4.4 minutes followed by a 99% B isocratic hold lasting 0.6 minutes (A: 0.1% aqueous formic acid, B: 0.06% formic acid in acetonitrile, 0.4 mL/min flow rate, 55 °C column temperature). Multiply-charged electrospray data (ESI+) were deconvoluted using the Waters’ MaxEnt 1 algorithm (Masslynx; Waters). The theoretical molecular weights of proteins encoding ncAAs were calculated by first computing the theoretical molecular weight of wild-type protein using an online tool (Expasy ProtParam; http://web.expasy.org/protparam/) and then manually corrected for the theoretical molecular weight of ncAAs following chromophore maturation.

### Macrocycle biosynthesis and purification

Chemically competent S3489 were transformed with plasmids encoding the synthetase, tRNA/qtRNA and desired macrocycle generating Npu intein. In all cases transformations were recovered for 2 h in 2xYT (United States Biological). All transformations were plated on 1.8% agar-2xYT plates (United States Biological) supplemented with relevant antibiotics: carbenicillin (50 µg mL^−1^), trimethoprim (10 µg mL^−1^) for two-plasmid circuits. The plates were incubated for 12–18 h in a 37 °C incubator. Colonies transformed with the appropriate EPs were picked the following day and grown in DRM containing carbenicillin (50 µg mL^−1^) and trimethoprim (10 µg mL^−1^). After growth for 16-24 hr at 37°C with 900 rpm shaking overnight cultures were diluted 200-fold in DRM with carbenicillin (50 µg mL^−1^), trimethoprim (10 µg mL^−1^), aTc (100 ng mL^−1^ or 15 ng mL^−1^ for triple suppression experiments), IPTG (1mM), L-Arabinose (10 mM), and chosen amino acid(s) (1mM). Following 16-24 h at 37°C with 900 rpm shaking, overnight cultures were centrifuged at 3000 x g for 5 min and supernatant discarded. Pellet was resuspended in 500 µL of PBS with 15% glycerol per mL of overnight culture, then centrifuged at 3000g for 5 minutes Supernatant was discarded, then the cell pellet was then vigorously vortexed and sonicated for 10 minutes, then subjected to three to five cycles of 10s liquid nitrogen, 2 min 42 °C, 2 min sonication. Following this, the pellet was resuspended in 50µL of acetonitrile and sonicated for 20-30 min. Finally, this was filtered through a 96-well PVDF filter plate (Corning^TM^ 3505) via centrifugation at 800 x g for 10 min. LCMS analysis was conducted on filtered samples by the Scripps Research Institute Automated Synthesis Facility, performed on a Waters I-Class LC and a Waters SQD2 single-quadrupole MS with a Waters Cortecs C18 column (1.6 um, 2.1×55 mm) using a 5-99% B gradient lasting 2.5 minutes (A: 0.1% aqueous formic acid, B: 0.06% formic acid in acetonitrile, 0.8 mL/min flow rate, 35 °C column temperature). Extracted ion chromatograms (XICs) and mass spectra were analyzed, collected, and exported using Waters Empower 3.7. Solid-phase peptide synthesis (SPPS) and purified macrocycle standards were generated by a commercial manufacturer (GenScript).

## Acknowledgements

The authors thank fellow Badran Lab members for their helpful discussions. The authors gratefully acknowledge Dr. Haoxin Li for assistance with NGS analyses; Maya L. Bulos for assistance with immunoblotting; Brian Seegers, Brian Monteverde, and Alphonse Owirka of the Scripps Research Flow Cytometry Core for their assistance with cell sorting; and Quynh Nguyen Wong, Brittany Sanchez, Jason Lee, and Dr. Jason Chen of the Scripps Research Institute Automated Synthesis Facility for their assistance with mass spectra analysis.

## Contributions

AC designed the study, lead experimental work, and analyzed results. AAP designed and led all macrocycle production studies. DLL contributed the orthogonal aaRS-tRNA matrix and codon– anticodon discovery efforts. ZL investigated reporter plasmid origin copy number changes to optimize quadruplet decoding. GDC designed the mRNA synonymous codon libraries. AHB conceived of and designed the study, performed experiments, analyzed results, and supervised the research. AC and AHB wrote the manuscript, with input from all authors.

## Data availability

The data found in Figures 1-6 and Extended Data Figures 1-27 are available in the associated Data Files 1-6. Select plasmids will be deposited to Addgene and NGS data will be uploaded to the NCBI SRA (SUB14446303).

